# Coupled feedback loops involving PAGE4, EMT and Notch signaling can give rise to non-genetic heterogeneity in prostate cancer cells

**DOI:** 10.1101/2020.12.29.423275

**Authors:** Divyoj Singh, Federico Bocci, Prakash Kulkarni, Mohit Kumar Jolly

**Author notes:** Authors to whom correspondence should be addressed (M.K.J.).

## Abstract

Non-genetic heterogeneity is emerging to be a crucial factor underlying therapy resistance in multiple cancers. However, the design principles of regulatory networks underlying non-genetic heterogeneity in cancer remain poorly understood. Here, we investigate the coupled dynamics of feedback loops involving a) oscillations in androgen receptor (AR) signaling mediated through an intrinsically disordered protein PAGE4, b) multistability in epithelial-mesenchymal transition (EMT), and c) Notch-Delta-Jagged signaling mediated cell-cell communication, each of which can generate non-genetic heterogeneity through multistability and/or oscillations. Our results show how different coupling strengths between AR and EMT signaling can lead to possible bistability in the levels of AR. These results reveal the emergent dynamics of coupled oscillatory and multi-stable systems and unravel mechanisms by which non-genetic heterogeneity in AR levels can be generated, which can act as a barrier to most existing therapies for prostate cancer patients.

## Introduction

Phenotypic plasticity, i.e. the ability of cells to switch back and forth reversibly among different states (phenotypes), is a universal feature of adaptation to varying environments encountered by various biological systems [1]. This theme has been investigated in developmental and evolutionary biology in detail [2,3], and is gaining importance in the context of disease progression as well [4–7]. Further, this theme is well-studied in cases of bacterial and yeast populations [8–12], and is increasingly being investigated for mammalian cells as well [13–15]. Mechanisms underlying phenotypic plasticity and consequent non-mutational or non-genetic heterogeneity, and their implications in determining the fitness of individual cells and entire cell populations remain to be comprehensively elucidated [16–18]. Recently, phenotypic plasticity has emerged as an important player in facilitating resistance against many standard chemotherapeutic drugs and targeted therapies for multiple cancers [19–21]. Similar to persisters in a bacterial population, drug-tolerant persisters have been observed in cancer [21]. Thus, besides the well-studied genetic/genomic heterogeneity in drug resistance, phenotypic (non-genetic) heterogeneity can enable adaptive cross-drug tolerance for cancer cells [22–24]. Unlike genomic changes that are ‘hard-wired’ and can be inherited by cell division, phenotypic changes are reversible and stochastic and may or may not be transmitted to the next generation [25–28].

Stochasticity in phenotypic plasticity and/or heterogeneity is a direct consequence of limited number effects of molecules involved in various biochemical reactions, including transcription, translation and cell-cell communication signaling [29–31]. Stochasticity can also drive cell-state transitions in a multistable system; this phenomenon may drive tumor repopulation after a phenotypic subpopulation has been selectively killed by the drug(s) [32]. Another regulatory level at which stochasticity can lead to cell-to-cell heterogeneity is promiscuity in protein-protein interactions. Such ‘conformational noise’ is manifested particularly by intrinsically disordered regions/proteins (IDRs/IDPs); many oncogenes/tumor suppressor genes have been shown to contain IDRs [33–36]. While both these modes of biological noise have been probed separately, their combined effect on cancer cell dynamics remains elusive. Here, we investigate the case of prostate cancer through the lens of these two mechanisms.

In prostate cancer (PCa), androgen-deprivation therapy (ADT) has been a standard-of-care treatment for over 75 years. Resistance to ADT eventually occurs in most patients, leading to metastatic castration-resistant prostate cancer (CRPC) [37]). Progression after ADT has been often connected to epithelial-mesenchymal transition (EMT) [38,39], a program that acts as the fulcrum of phenotypic plasticity. Multistability in EMT, driven by mutually inhibitory feedback loops among transcription factors and microRNAs [40], can lead to transitions among multiple cell states – epithelial (E), mesenchymal (M) and hybrid E/M – as observed for prostate cancer cells [41]. On the other hand, intrinsic disorder in the cancer/testis antigen PAGE4, has been suggested to regulate the signaling of androgen receptor (AR), a target of ADT. Differently phosphorylated versions of PAGE4 can form a negative feedback loop involving AR, which has been predicted to give rise to oscillations [42,43], thereby generating non-genetic heterogeneity in the levels of AR in an isogenic population. AR can form a mutually inhibitory loop with ZEB1, an EMT-inducer [38], thus coupling a multistable system with an oscillatory one. The emergent dynamics of such coupling, particularly in the context of PCa, is unclear. Therefore, here we simulate the coupled dynamics of AR-EMT circuit and demonstrate that depending on the strength of bidirectional coupling between ZEB1 and AR, multistability can be seen in AR signaling and/or some oscillations can be seen in EMT circuitry. Besides intracellular signaling, we also investigate the effect of this coupling on Notch signaling, a cell-cell communication pathway. Our results reveal how AR signaling can display different nonlinear emergent dynamics (oscillations, bistable) and therefore generate and maintain non-genetic heterogeneity in an isogenic cancer cell population.

## Results

### Oscillations and bistability in PAGE4-AR signaling

We have previously investigated standalone dynamics of PAGE4-AR and EMT circuits [43,44]. The PAGE4-AR circuit consists of three relevant PAGE4 phospho-forms (WT-PAGE4, HIPK1-PAGE4, CLK2-PAGE4), together with c-Jun and androgen receptor (AR). Because c-Jun potentiation via HIPK1-PAGE4 can eventually drive the hyper-phosphorylation of HIPK1-PAGE4 to a short-lived CLK2-PAGE4, a delayed negative feedback loop is formed that can drive oscillations in this circuit. This hyperphosphorylation of HIPK1-PAGE4 to CLK2-PAGE4 involves AR which is inhibited by c-Jun potentiation and can inhibit CLK2 upregulation. The strength of this negative feedback is quantified by a fold-change parameter (*λ*_*PAGE*4_), so that *λ_PAGE4_* = 0 indicates maximal inhibition on AR (by c-Jun) and CLK2 (by AR). On the other hand, *λ*_*PAGE*4_ = 1 indicates no inhibition for either of the two links at all, thus, the feedback loop is broken (*see Methods*). Depending on this feedback strength (value of *λ*_*PAGE*4_), AR levels can either display sustained oscillations (strong inhibition, **Fig 1A, i**) or relax to a steady level (weak inhibition, **Fig 1A, ii**).

**Figure 1:**
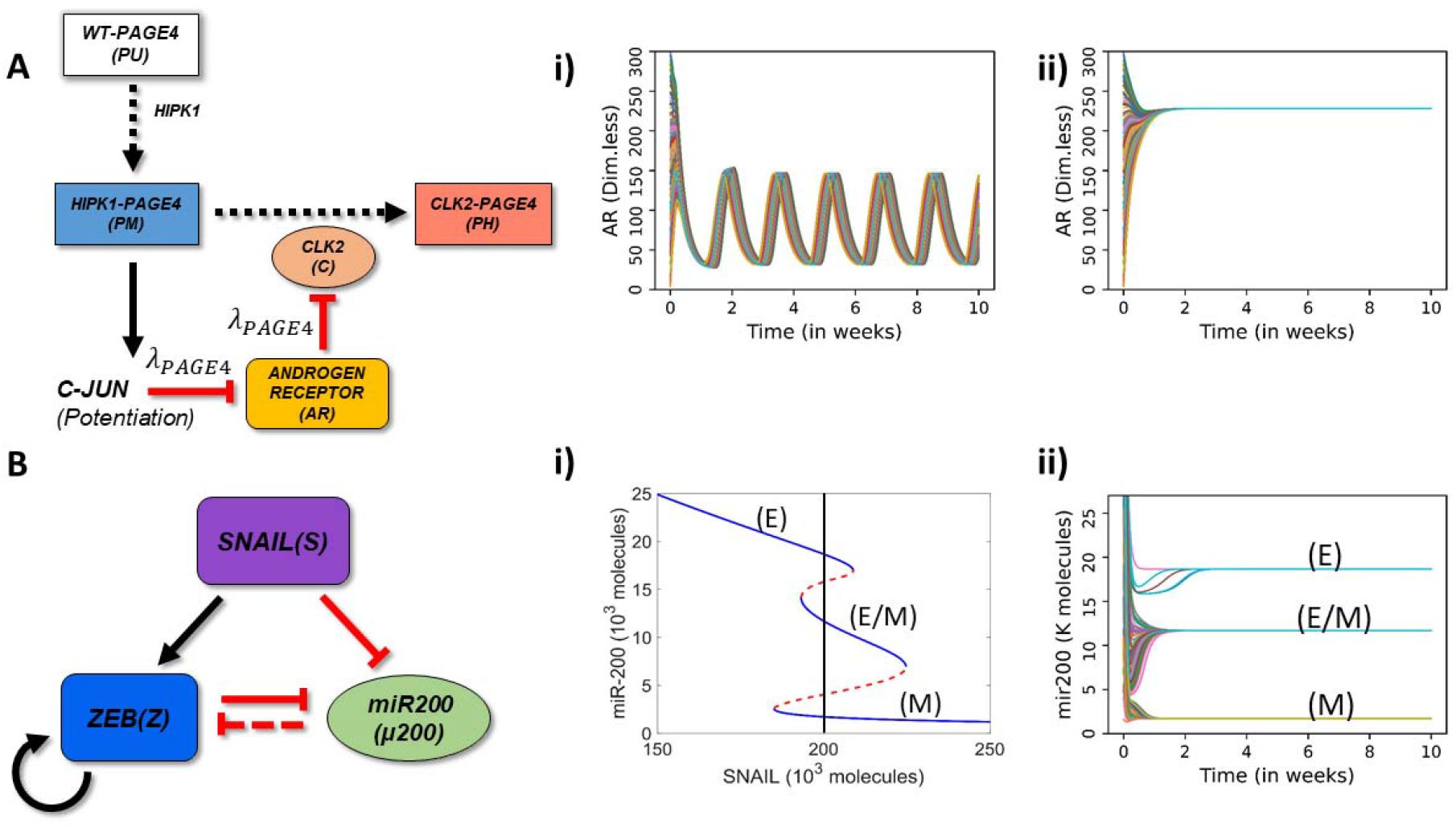
Schematic representation of PAGE4-AR and EMT circuits and their standalone dynamics. ***A)** Schematic representation of PAGE4-Androgen Receptor (AR) circuit: The enzyme HIPK1 double phosphorylates WT-PAGE4 and forms the HIPK1-PAGE4 complex which can be further hyper-phosphorylated by CLK2 enzyme. Solid arrows show activation, dotted arrows show phosphorylation and red hammer heads show inhibition. In turn, the HIPK1-PAGE4 complex regulates CLK2 levels via the intermediates c-JUN and AR. A strong inhibition of AR by c-JUN and that of CLK2 by AR leads to oscillations (λ*_*PAGE*4_ = 0.1*) (i) or a single steady state (mono-stability) (λ*_*PAGE*4_ = 0.9*) (ii). **B)** EMT circuit: ZEB and microRNA-200 form a mutually inhibiting loop while SNAIL acts as an external EMT inducer. Solid arrows show transcriptional activation, dashed line show microRNA-mediated inhibition, and solid hammerheads show transcriptional inhibition. (i) Bifurcation diagram of microRNA (miR)-200 as a function of SNAIL shows tristability, bistability or mono-stability depending on SNAIL levels. Blue and red curves show stable and unstable states respectively. The vertical black line depicts the SNAIL level (=200,000 molecules) used in panel ii. ii) Dynamics of miR-200 for SNAIL = 200K showing the existence of three states -epithelial (high miR-200; 20K molecules), mesenchymal (low miR-200; < 2K molecules) and hybrid E/M (medium miR-200; ~12K molecules). In panels Ai-ii, Bii, different curves depict AR and miR-200 dynamics starting from multiple randomized initial conditions*.

Next, in the core EMT circuit, ZEB and miR-200 inhibit each other, and ZEB can self-activate. SNAIL acts as an external signal driving EMT by activating ZEB and inhibiting miR-200 [44]. Thus, a bifurcation diagram of miR-200 levels with respect to SNAIL displays transition from an epithelial (high miR-200, low ZEB) to a hybrid epithelial/mesenchymal (medium miR-200, medium ZEB) to a mesenchymal (low miR-200, high ZEB) state. Therefore, the standalone SNAIL/ZEB/miR-200 circuit can behave as a monostable, bistable, or tristable system with coexistence of E, E/M and M phenotypes depending on the level of SNAIL-driven EMT induction (**Fig 1B, i-ii**).

Before coupling the PAGE4-AR circuit with the EMT circuit, we first add a single node X to the PAGE4-AR circuit to understand how perturbations to AR signaling modify its oscillatory dynamics **(Fig 2A)**. We also allow X to self-inhibit or self-activate. This coupling between AR and X mimics the scenario of mutual inhibition between AR and ZEB1, and possible self-activation of ZEB1 [45]. We investigated the dynamics of this extended circuit at varied strengths of coupling between AR and X (represented by *λ_DNFL_*), and different strengths of interaction in the PAGE4-AR feedback loop (represented by *λ*_*PAGE*4_). These values vary between 0 and 1, and the smaller the value, the stronger the inhibition (see Methods). In this circuit, *λ_DNFL_* represents the fold-change (and hence the strength of regulation) for both links: the inhibition of AR by X and *vice versa*.

**Figure 2:**
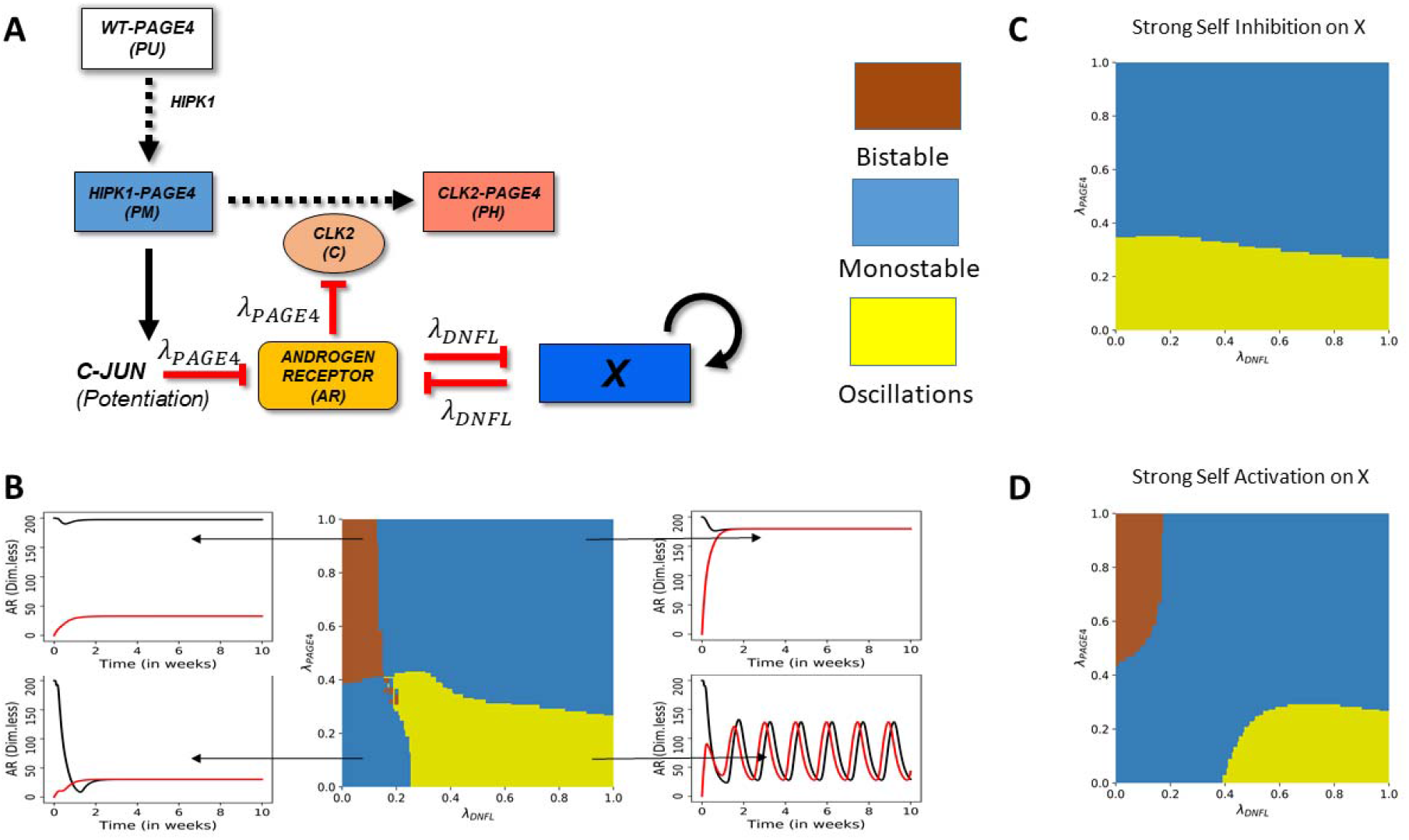
Perturbation of AR signaling can lead to monostable, bistable, or oscillatory dynamics. ***A)** Circuit showing the connections of a generic node (X) with the PAGE4-AR circuit. X can also transcriptionally regulate itself (blue arrow). **B)** Phase diagram of the PAGE4-X circuit as a function of strength of PAGE4 internal coupling λ_PAGE4_) and that of double negative feedback loop coupling of AR with X (λ_DNFL_). Self-regulation of X is ignored here. Inset panels show representative examples of dynamics in different phases, such as bistability (top left), mono-stability leading to low AR levels (bottom left), monostability leading to high AR levels (top right), and oscillations (bottom right). Black and red curves indicate two different initial conditions of high AR and low AR respectively. **C)** Same as B) for a strong self-inhibition of X: (λ_XtoX_* = 0.1) ***D)** Same as B) for a strong self-activation of X: (λ_XtoX_ = 7.5). In Panels B-C-D, yellow shading indicates oscillations, blue indicates monostability, and brown indicates bistability*.

At high *λ*_*PAGE*4_ and low *λ_DNFL_*, i.e. when AR and X inhibit each other strongly but the internal interactions in the PAGE4-AR circuit are weak, the system shows a bistable behavior – coexistence of (high AR, low X) and (low AR, high X) states, typical of double negative feedback loops [46,47] (**Fig 2B; center panel**, top left region). Conversely, at low *λ*_*PAGE*4_ and high *λ_DNFL_*, i.e. when AR and X inhibit each other moderately but PAGE4-AR circuit features a strong negative feedback loop, the system displays sustained oscillations, typical of delayed negative feedback loops [48–50] (**Fig 2B**; center panel, bottom right region). Interestingly, at both (high *λ*_*PAGE*4_, high *λ_DNFL_*) and (low *λ*_*PAGE*4_, low *λ_DNFL_*), AR is mono-stable, albeit at different steady state levels (**Fig 2B**, center panel; top right and bottom left regions). In the former case, AR saturates at a higher level, perhaps due to lack of inhibition by X and/or other components of PAGE4-AR circuit. Conversely, in the latter case, AR saturates to a lower level. Therefore, this coupled circuit can show bistable, monostable or oscillatory dynamics depending on relative strengths of the links.

Next, we study the effects of self-regulation on X, denoted by the fold-change parameter *λ_XtoX_*. A value greater than 1 implies self-activation of X, whereas a value smaller than 1 implies self-inhibition of X. In the case of self-inhibition (*λ_XtoX_ =* 0.1), the bistable phase disappears and the system exhibits either monostable dynamics or oscillations (**Fig 2C**). This trend is consistent with observations that positive feedback loops facilitate multistability [40]. Oscillations are noted only at strong coupling within the PAGE4-AR feedback loop (i.e. smaller values of *λ*_*PAGE*4_) (**Fig 2C**, yellow-shaded region); for weak PAGE4-AR coupling (i.e. higher values of *λ*_*PAGE*4_), AR saturates at high steady state values, irrespective of the coupling strength between AR and X (**Fig 2C**, blue-shaded region). Conversely, in case of self-activation of X, the system largely behaves as in the case of no-self regulation, although with an increased parameter regime for bistability (**Fig 2D**).

To further gain confidence in these observations, we investigated the effect of altering other model parameters. All non-phosphorylation reactions here are modeled via shifted Hill function which describe the production fold-change of a given species as a function of inducer/inhibitor levels (see Methods); specifically, a Hill coefficient (n) quantifies how nonlinearly the foldchange depends on inducer/inhibitor level. We observed that the higher the value of n for AR-X coupling (*n_DNFL_*) and the lower the value of n for coupling within PAGE4-AR circuit (*n_PAGE4_*), the larger the parameter region enabling bistability. Conversely, lower values of *n_DNFL_* and/or higher values of *n*_*PAGE*4_ drives oscillations **(Fig S1A).** Further, we have so far considered identical parameters for the shifted Hill functions denoting the inhibition of AR by c-Jun and that of CLK2 by AR. For the purpose of sensitivity analysis, even when we considered them to be independent parameters, we noticed that both these links have to be strong (i.e. *λ*~0) to facilitate oscillations in the levels of AR (**Fig S1B**). Put together, all these results suggest that a strong ‘external coupling’ (i.e. double negative feedback loop between AR and X) favors bistability, while a strong ‘internal coupling’ (i.e. negative feedback loop formed between AR and phospho-forms of PAGE4) drives oscillations in PAGE4-AR-X coupled circuit.

### Dynamics of coupled PAGE4-AR-EMT circuits

Next, we investigate the dynamics of coupled PAGE4-AR and EMT circuit. In this coupled circuit, AR and ZEB1 inhibit each other, and Snail (S) is an external EMT-inducer **(Fig 3A)**. To investigate how the AR-ZEB coupling modifies the dynamics of the coupled circuit, we probe the system’s dynamics for various strength combinations of the AR-to-ZEB inhibition (described by the fold-change *λ_AtoZ_*) and ZEB-to-AR inhibition (described by the fold-change *λ_ZtoA_*). Given that the coupled PAGE4-EMT circuit includes time delay and can potentially give rise to oscillations, we inspect the behavior of the circuit by evaluating its temporal dynamics for various combinations of coupling strengths and SNAIL level. As a first step, we study the effects of weak bidirectional coupling, i.e. *λ_AtoZ_* = *λ_ZtoA_*= 0.9. As expected, here, both circuits largely show their standalone dynamics, *i.e*. oscillations for PAGE4-AR circuit and monostability/multistability for the EMT circuit depending on the SNAIL levels. Interestingly, the EMT circuit can show oscillations of very small magnitude on top of its steady states obtained, especially at SNAIL levels enabling multistability **(Fig S2)**.

**Figure 3:**
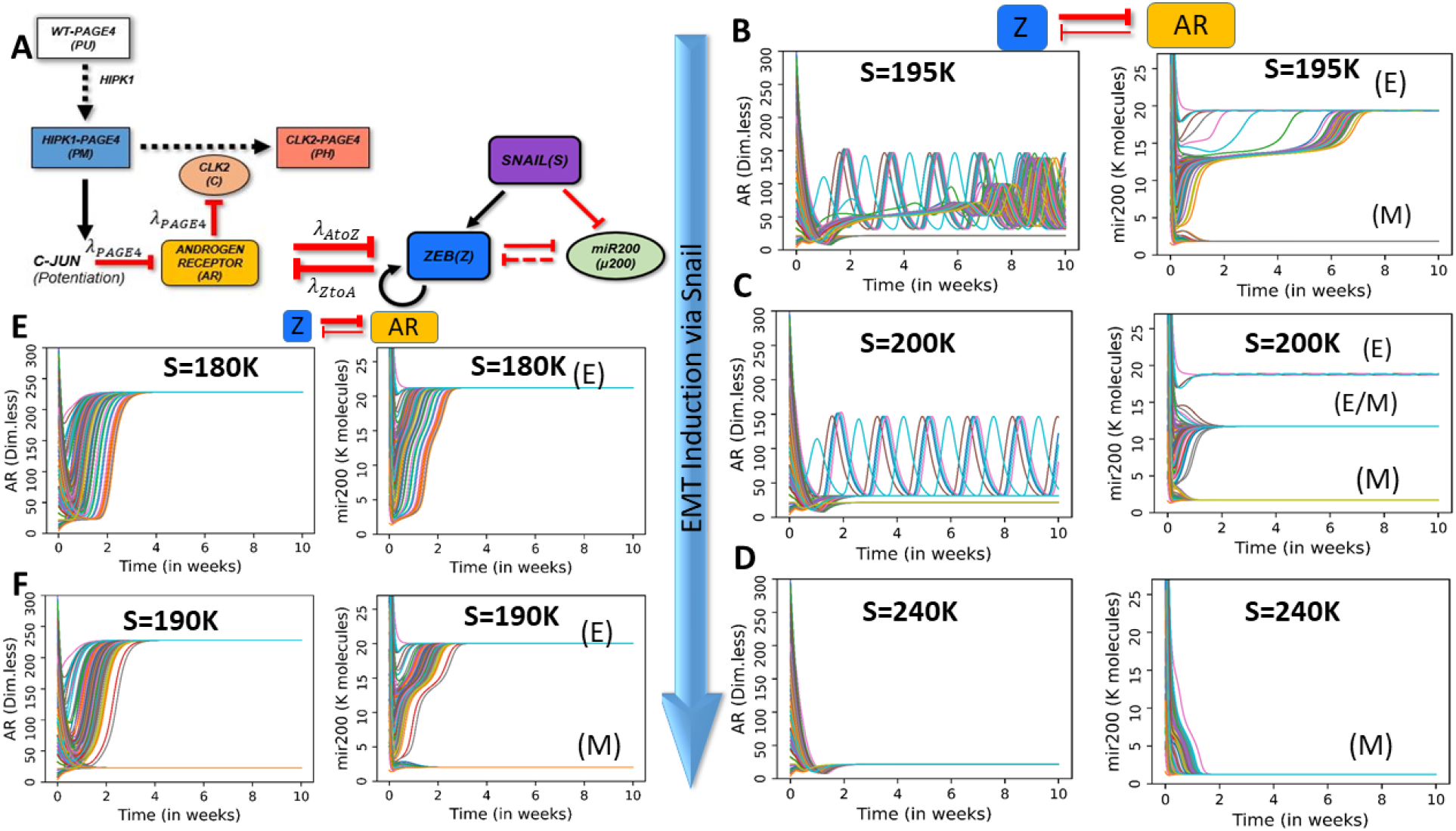
Dynamics of PAGE4-AR-EMT coupling for case of strong inhibition of AR by ZEB1. ***A)** Coupled PAGE4-AR and EMT networks. **B-F)** Dynamic trajectories of AR and miR-200 for strong inhibition of AR by ZEB1, but weak inhibition of ZEB1 by AR (λ_AtoZ_* = 0.9, *λ_ZtoA_* = 0.1). ***B-D)** Dynamics at different Snail (S) values for strong internal coupling in PAGE4-AR circuit (λ*_*PAGE*4_ = 0.1). ***E-F)** Same as B-D, but for weak internal coupling (λ_AtoZ_* = 0.9). *Dynamic trajectories are plotted for 100 different initial conditions for a period of 10 weeks. Corresponding values of SNAIL are mentioned in respective panels*.

Next, we increase the coupling strength on one arm: *λ_ZtoA_* = 0.1 (ZEB inhibits AR strongly). For low Snail values (S=160K), AR oscillates whereas miR-200 relaxes to a high value typical of the epithelial state **(Fig S3A)**. As Snail levels are increased (S=195K), the circuit can attain two possible stable states – an epithelial (high miR-200) state with AR oscillations or a mesenchymal (low miR-200) state with low AR (**Fig 3B**). On further increasing Snail levels (S=200K), a third hybrid E/M (medium miR-200) state emerges. **(Fig 3C)**. While the epithelial state allows AR oscillations, a partial or complete EMT (i.e. hybrid E/M or M state) tends to suppress oscillations. Further increase of Snail levels (S=215 K) eliminates the epithelial state as expected, and the EMT circuitry shows coexistence of the hybrid E/M and mesenchymal states **(Fig S3B)**. An even higher value of Snail (S=240K) drives a monostable mesenchymal phase in the EMT bifurcation diagram (**Fig 1B**). For these values of Snail, oscillations in AR disappear and the AR dynamics converge to low steady state values, given the higher levels of ZEB and consequently a strong inhibition of AR by ZEB (**Fig 3C, D**). Overall, in the case of strong inhibition of AR by ZEB, the multistability of the EMT circuit can be propagated to PAGE4-AR circuit, and SNAIL-driven EMT induction can dampen the PAGE4-AR oscillations.

The above-mentioned results consider a case of strong PAGE4 ‘internal coupling’, i.e. in the parameter region where the standalone dynamics of PAGE4-AR circuit is oscillatory (*λ*_*PAGE*4_ = 0.1). Conversely, in the case of weak internal coupling (*λ*_*PAGE*4_ = 0.9), AR saturates to a high steady state value (monostable) at low Snail values (S=180K) **(Fig 3E)**. Increasing Snail levels (S=190K), however, induces bistability in AR levels, such that depending on the initial condition, cells can converge to either an (epithelial, high AR) state or a (mesenchymal, low AR) state **(Fig 3F)**. Thus, multistability is passed on from EMT circuit to the PAGE4-AR circuit in this case too.

Next, we examine the case when we interchange the interaction strengths of both arms, i.e. when AR inhibits ZEB1 strongly, but ZEB1 inhibits AR weakly (*λ_AtoZ_* = 0.1, *λ_ZtoA_* = 0.9). Due to the weaker effect of ZEB1 on AR, AR oscillates for any value of SNAIL (**Fig 4A,i; S3C**.). On the other hand, the dynamics of the EMT circuit is largely driven by AR. Interestingly, miR-200 does not show multistability, instead it oscillates, and the amplitude of oscillations increases with increasing values of SNAIL (S=240K and S=300K) (**Fig 4A, ii-iv**).

**Figure 4:**
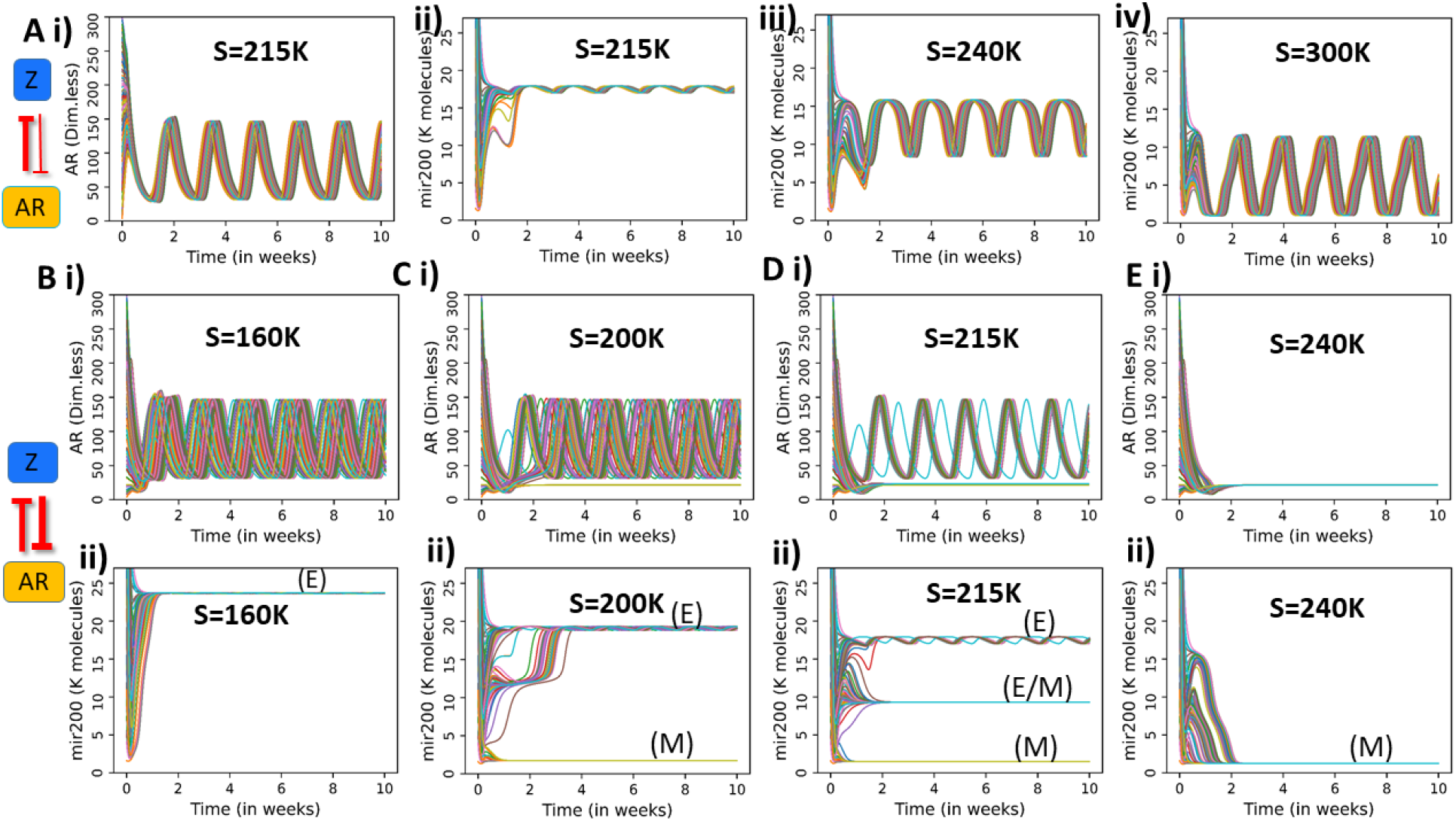
Dynamics of PAGE4-AR-EMT coupling, for case of strong inhibition of ZEB1 by AR. ***A)** Dynamic trajectories of AR and miR-200 for the case of strong effect of AR on ZEB1 but weak effect of ZEB1 on AR (λ_AtoZ_=0.1 and λ_ZtoA_=0.9). **B-E)** Dynamic Trajectories of AR (B) and miR-200 for the case of strong effect of AR on ZEB1 and strong effect of ZEB1 on AR (λ_AtoZ_=0.1 and λ_ZtoA_=0.1).*

Finally, we consider the case of strong coupling on both arms, i.e. when AR and ZEB1 both inhibit each other strongly (*λ_AtoZ_* = *λ_ZtoA_*= 0.1). For low Snail values (S=160K), AR shows oscillations and miR-200 assumes an epithelial level (**Fig 4B**). However, at increasing Snail values (S=200K, S=215K), AR shows the coexistence of an oscillatory state and steady state corresponding to low AR values (**Fig 4C,D**). Corresponding to these cases, miR-200 can either assume an epithelial level with small oscillations, or a hybrid E/M or mesenchymal state without any oscillations. Upon further increasing SNAIL levels (S=240K), only the mesenchymal state exists in terms of EMT, and correspondingly AR saturates to a low steady state value, driven by both high levels of ZEB1 and strong inhibitory effect of ZEB1 on AR considered here (**Fig 4E**).

### Integrating Notch-Delta-Jagged signaling circuit with combined PAGE4-EMT circuit

So far, we have considered phenotypic heterogeneity emerging from intracellular signalling. Communication between cells, however, can potentially modulate the PAGE4-EMT dynamics and propagate heterogeneity to other cells. Specifically, Notch signalling is widely recognized as a crucial mediator of cancer progression that operates via ligand-receptor binding between neighbouring cells [51]. Previously, we have studied the coupled dynamics of EMT and Notch that were capable of exhibiting a maximum of four distinct states [52]. Drawing from these studies, here, we combined the three networks (PAGE4-AR, EMT, Notch-Delta-Jagged) at a single-cell level. Notch signaling is activated by transactivation of Notch receptor by Delta and/or Jagged ligands, thus we investigate the dynamical behavior of the coupled circuit at varied levels of such ligands. In other words, we model the coupled PAGE4-EMT-Notch circuit in an individual cell that is exposed to an varying external levels of Notch receptors (Notch) and ligands (Delta, Jagged). These external receptors and ligands mimic the effect of neighbouring cells and activate the intracellular Notch signalling cascade. Increasing the levels of external Delta ligand (D_ext_) can drive EMT through Notch signaling mediated activation of SNAIL, subsequently affecting the dynamics of EMT and PAGE4-AR circuits. We have previously shown that the dynamics of EMT circuit remain largely unaltered upon including miR-34 [44]; thus, we included miR-34 in our circuit, given its connections with Notch signaling circuit [52] (**Fig 5A)**. The EMT-Notch circuit (i.e. without incorporating the PAGE4-AR circuit) can assume up to four distinct phenotypes when induced with varying levels of external Delta ligand (D_ext_) **(Fig 5B, Fig S4).** In particular, when comparing to the standalone EMT bifurcation (see **Fig 1B,i**), we notice that the epithelial branch (high miR-200) splits into two distinct branches in the EMT-Notch coupled circuit: a (high miR-200, high Notch), i.e. epithelial Receiver state, and a (high miR-200, high Delta), i.e. epithelial Sender state. On the other hand, both the hybrid E/M (medium miR-200) and M (low miR-200) branches are characterized by a Sender/Receiver Notch state with (high Notch, high Jagged).

**Figure 5:**
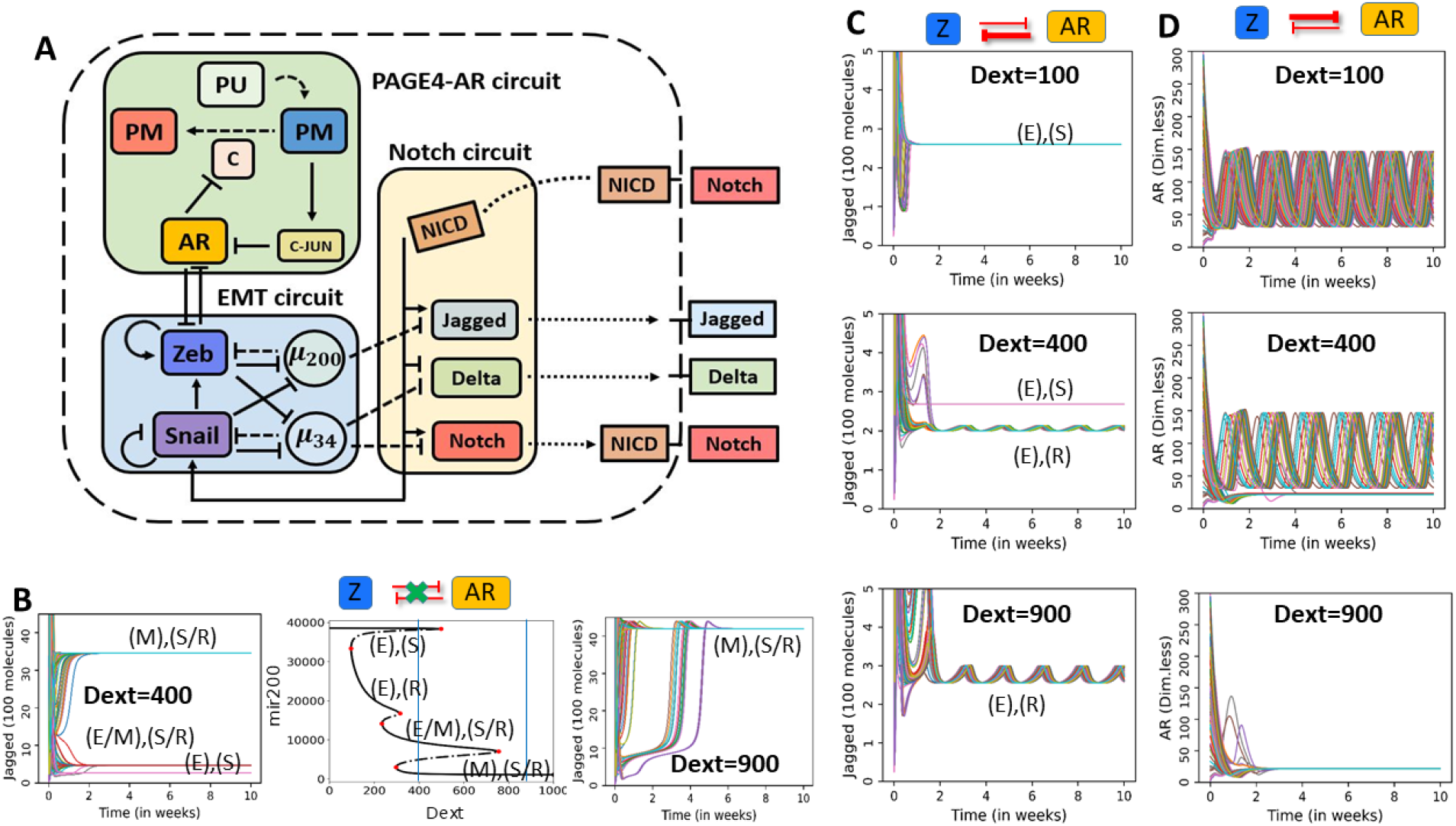
Dynamics of coupled EMT, PAGE4-AR and Notch signaling: ***A)** Schematic of the coupled circuit. **B)** Central panel: bifurcation diagram of miR-200 as a function of external Delta ligands (Dext) in Notch-EMT circuit (i.e. no coupling with the AR-PAGE4 circuit). Left and right panels show temporal dynamics of Jagged for two values of Dext (highlighted by vertical dotted lines in central panel). **C)** Temporal dynamics of AR for strong AR-to-ZEB signalling and weak ZEB-to-AR signalling (λ_AtoZ_ =0.1 and λ_ZtoA_=0.9), for increasing values of external Delta ligands (Dext). **D)** Same as C) but for λ_AtoZ_=0.9 and λ_ZtoA_ =0.1. In panels B-D, different colours depict trajectories starting from distinct random initial conditions. Unit of Dext is number of molecules.*

We first consider the case of strong AR-to-ZEB signalling and weak ZEB-to-AR signalling (*λ_AtoZ_* = 0.1, *λ_ZtoA_* = 0.9), thus studying how AR dynamics propagates to EMT and Notch **(Fig 5C, S5A)**. For a low value of D_ext_ (D_ext_ =100), the cell assumes an epithelial phenotype, and Notch receptors and ligands relax to a constant level (**Fig 5C**, top panel). A higher D_ext_ level (D_ext_ =400) induces a co-existence of two epithelial phenotypes – one of them (epithelial Sender) does not exhibit oscillations in terms of miR-200 and Jagged and the other one (epithelial Receiver) does (**Fig 5C**, middle panel; S5A). On further increasing Dext (D_ext_ =900), the epithelial Sender phenotype is not observed, perhaps because of strong activation of Notch signaling and consequent inhibition of Delta by Notch Intra Cellular Domain (NICD). In this scenario, the epithelial Receiver phenotype is maintained which continues to exhibit oscillations in both miR-200 and Jagged (**Fig 5C**, bottom panel; **S5A**). Overall, a stronger activation of Notch signaling can drive propagation of oscillations seen in PAGE4-AR circuit to the Notch circuit as well through the intermediary EMT circuit (please note that Notch circuit and PAGE4-AR circuits are connected solely through the EMT circuit here).

Next, we consider the opposite case, i.e., weak AR-to-ZEB signalling and strong ZEB-to-AR signalling (*λ_AtoZ_* = 0.9, *λ_ZtoA_* = 0.1) and study how Notch multistability affects AR oscillations **(Fig 5D, S5B)**. At low values of Dext (Dext=100), all initial conditions converge to an epithelial phenotype where ZEB levels are low and thus cannot inhibit AR oscillations (**Fig 5D**, top panel). At intermediate Dext (Dext=400), AR can either maintain its oscillatory behaviour or relax to a low steady state level (**Fig 5D**, middle panel). On further increasing Dext (Dext=900), AR loses its oscillations and saturates at a low steady state (**Fig 5D**, bottom panel). Thus, in case when ZEB inhibits AR strongly, activation of Notch signaling can dampen AR oscillations through increased ZEB levels.

In case of strong inhibition on both sides (*λ_AtoZ_* = 0.1, *λ_ZtoA_* = 0.1), we see trends reminiscent of both scenarios discussed above. At low Dext value (Dext=100), epithelial Sender phenotype is observed, and AR shows oscillations (**Fig S6A**). As Dext values are increased, Notch signaling is activated that can increase the levels of SNAIL to a value that supports multistability in EMT. Thus, at Dext=400, we observe 4 states in coupled EMT-Notch-PAGE4 circuit as seen previously – epithelial Sender ((E), (S)), epithelial Receiver ((E), (R)), hybrid E/M Sender/Receiver ((E/M), (S/R)) and Mesenchymal Sender/Receiver ((M), (S/R)) (**Fig S6B**). Concurrently, AR dynamics shows the co-existence of oscillations and a low steady state value. On increasing Dext further (Dext=900), the epithelial sender ((E), (S)) and hybrid E/M Sender/Receiver ((E/M), (S/R)) states are not observed, consistent with trends expected of a stronger activation of Notch (**Fig S6C).**

## Discussion

Non-genetic heterogeneity is a central theme in cellular decision-making in embryonic development [53]. Its importance in cancer is beginning to be appreciated, with an increasing quantitative and systems level analysis of such heterogeneity to understand the mechanistic underpinnings [54]. Here, we investigate the coupled dynamics of three signaling networks, each of which is capable of generating non-genetic heterogeneity in a cancer cell population. Two of these circuits (EMT, Notch-Delta-Jagged) are multistable [55–58], and the third one (PAGE4-AR) can exhibit oscillations [42,43]. In our previous work, we had exemplified how the coupled dynamics of both multistable modules (EMT and Notch-Delta-Jagged signaling) at a multi-cell level drove varied spatiotemporal patterns of E, M and hybrid E/M phenotypes [59]. Here, we interrogate the emergent dynamics of coupling of circuits exhibiting multistable and oscillatory behavior and discuss its implications for prostate cancer cells.

Depending on relative strengths of the effect of ZEB1 on AR and *vice-versa*, we observed that the stand-alone dynamical features of EMT and AR circuits – multistability and oscillations – could percolate to the other circuit, i.e. EMT circuit may show oscillations and/or AR circuits may exhibit bistability. While many features of multistability in EMT has been experimentally observed in multiple cancer cell lines (co-existence of multiple states [60], hysteresis [61,62], and spontaneous state switching [41]), oscillations in EMT have not yet been observed. Encouraging results from preliminary studies using GFP-AR to quantify nuclear translocation of AR in individual PCa cells [63] suggests that empirically validating our theoretical results is technically feasible.

The bidirectional coupling between AR signaling and EMT offer a possible mechanistic link associating the progression of cells towards a partial or full EMT and gain of therapy resistance in prostate cancer. In particular, our model suggests that the epithelial phenotype usually cooccurs with PAGE4 oscillations, whereas transitions to hybrid E/M or mesenchymal phenotypes quench these oscillations and promote low AR levels. From a clinical standpoint, low levels of AR suggest an androgen independent (AI) phenotype that is potentially less susceptible to androgen deprivation therapies. Therefore, EMT induction can potentially promote therapy resistance by stabilizing an androgen independent PCa phenotype through the ZEB1-AR signaling axis. A partial and/or full EMT has been associated with resistance to various therapies in multiple cancers [64]; however, a comparative analysis of partial and full EMT in terms of therapeutic resistance is beyond the scope of the model considered here. Nonetheless, such AR-ZEB1 coupling suggests that not only EMT can drive the acquisition of a drug-resistant state as observed *in vitro* and *in vivo* [65,66], but conversely, a switch from drug-sensitive to drug-resistant state can also trigger EMT. Residual breast cancers after conventional therapies have been shown to be mesenchymal [67], but this analysis was at a bulk population level. Thus, these results preclude us from discerning whether the residual cells are a result of phenotypic switching or they are derived from pre-existing mesenchymal cells that were selected during the therapy.

Our model predicts that besides the EMT circuit, Notch signaling pathway can also exhibit oscillations. This pathway has been shown to display oscillatory dynamics, but mostly in the context of somite segmentation clock [51]. The oscillations predicted by our model for EMT and Notch signaling is predicated on a) oscillations in AR levels that remain to be experimentally tested, and b) the mutual inhibition between ZEB1 and AR. *In vivo* imaging has revealed oscillatory dynamics of ER (estrogen receptor) transcriptional activity in a tissue-dependent manner [68]. Negative feedback loops such as p53-MDM2 can give rise to oscillations [48]; whether such a loop exists for ER and AR (in addition to the one including PAGE4) in PCa cells, remains to be determined. Furthermore, the mutual inhibition between ZEB1 and AR may only be true for PCa cells, but not for other cancers [69]. Finally, any direct coupling between PAGE4/AR and Notch-Delta-Jagged signaling can alter the emergent dynamics shown for these circuits.

Considered together, our work highlights how coupling between Notch, EMT and PAGE4/ AR signaling pathways can give rise to non-trivial dynamics and non-genetic heterogeneity in a cancer cell population. These results are likely to impact how we treat PCa in the future.

## Data Availability

All codes are available publicly on the GitHub page of DS (https://github.com/Divyoj-Singh/PAGE4-AR-EMT-NDJ)

## Author contributions

DS and FB performed research; DS, FB and PK analyzed data; MKJ supervised and conceived research; all authors contributed to writing and editing of manuscript.

## Materials and Methods

### Mathematical Modelling of the coupled PAGE4/AR-EMT-Notch circuit

In our model, the temporal dynamics of micro-RNAs or proteins (say, X) obey the chemical rate equations that describe their interactions with other species in the circuit. The generic form of such equations is:

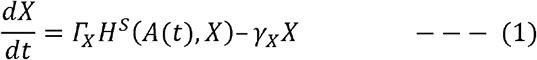

Here, the first term in RHS (*Γ_X_H^S^*(*A*(*t*), *X*)), stands for the net production rate of X. This production rate depends on the basal production rate of X (*Γ_X_*) and a function *H^S^*(*A*(*t*), *X*) which represents the regulation of levels of X resulting from interactions with any another species in the circuit (say, A). In case of regulation on X from multiple species, those terms are multiplied. The second term on the RHS (−*γ_X_X*), represents the first-order degradation of X. We used Shifted Hill function to model the effect of one species on another, which leads equation (1) to take the following form:

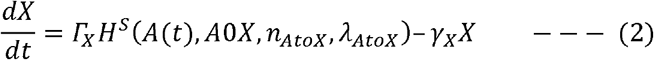

Where H^S^ takes the following form:

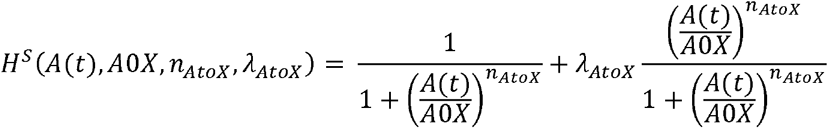

The first argument in the Shifted Hill function (A(t)) is number of molecules of the species A at any given time (t). Furthermore, in the PAGE4 circuit, we consider that case of delay in the inhibition of CLK2 by AR. In this particular case, the levels of CLK2 at time t depend on the levels of AR at (*t* - *τ*), where *τ* expresses the amount of delay. A0X is the half-maximal concentration parameter of the Hill function. *λ_AtoX_* is the fold-change of X due to the effect of A. *λ_AtoX_ >* 1 corresponds to activation, and a higher value of *λ_AtoX_* implies a stronger activation. Conversely, *λ_AtoX_* < 1 corresponds to inhibition, and the closer the value of *λ_AtoX_* to 0, the stronger the inhibition strength. *λ_AtoX_* = 1 corresponds to the limit case where X does not affect A. Additionally, *n_AtoX_* is the characteristic Hill function coefficient which determines the steepness of the Hill function.

Detailed equations for all species in the PAGE4, EMT and Notch circuits are presented in the Supplementary Information (Section 1-3).

