## Supplementary material for "Coupled feedback loops involving PAGE4, EMT and Notch signaling can give rise to non-genetic heterogeneity in prostate cancer cells": SI Figures

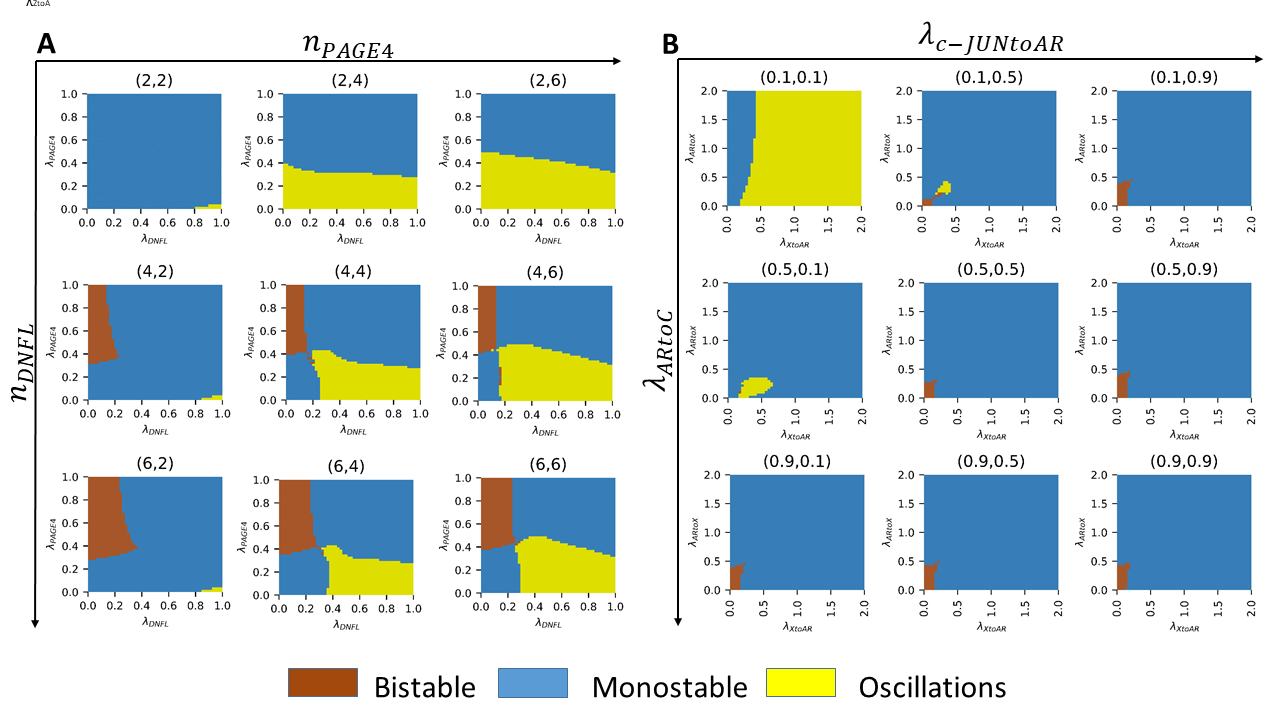

***Figure S1: Sensitivity analysis*** ***A)*** *Effect of Hill-coefficients:* *Phase plot between* $\lambda_{DNFL}$ *and* $\lambda_{PAGE4}$*for different values of Hill-coefficients for both these couplings. Yellow shading indicates oscillations, blue indicates monostability, and brown indicates bistability.* ***B)*** *Effect of different coupling strengths:* *Phase plot between* $\lambda_{AtoX}$ *and* $\lambda_{ZtoA}$ *for different strengths of internal coupling interactions (λ HIPK1-PAGE4 complex to AR (*$\lambda_{PMtoA}$*) and Lambda AR to CLK2 (*$\lambda_{AtoC}$*)). For strong internal coupling (0.1, 0.1), the system shows oscillations even when X and A activate each other (*$\lambda> 1$*). But when the internal coupling is weak (0.9, 0.9), the system is largely monostable mainly and bistable in a small parameter region.*

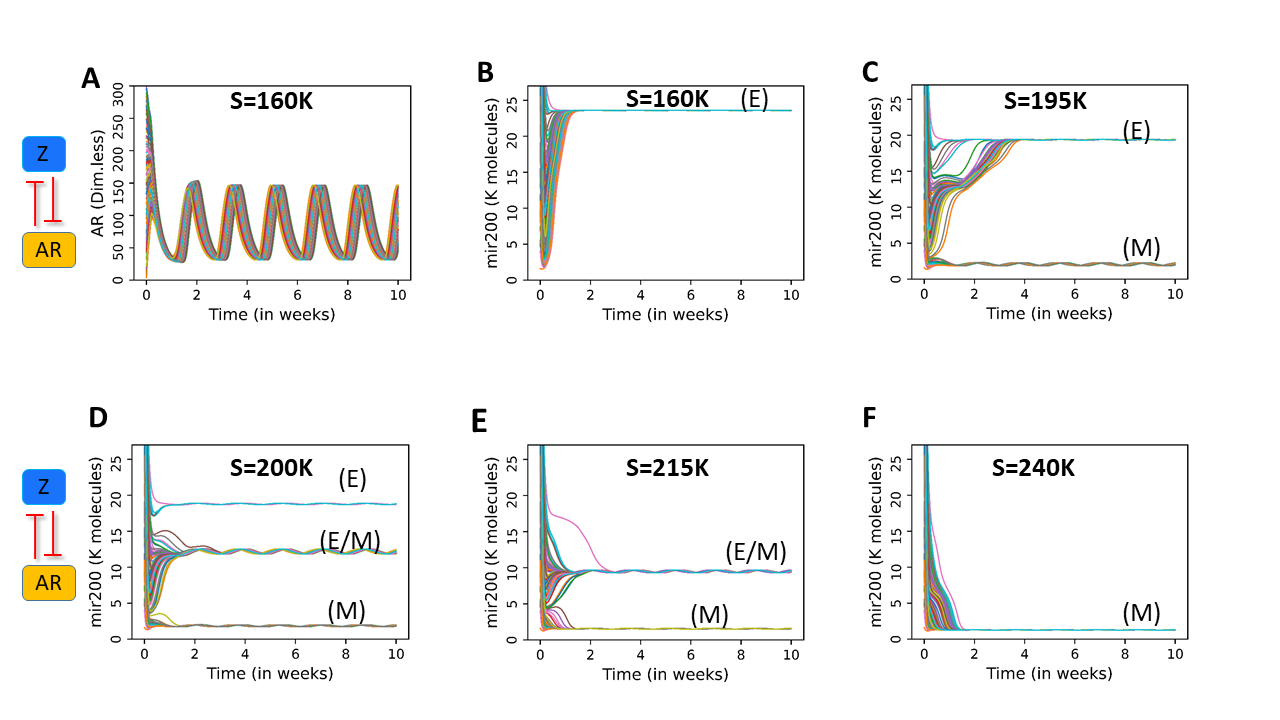
***Figure S2: Dynamics at weak coupling between the EMT and PAGE4/AR circuits (***$\boldsymbol{\lambda}_{\boldsymbol{AtoZ}}$ ***and*** $\boldsymbol{\lambda}_{\boldsymbol{ZtoA}}$ ***=0.9):*** ***A)*** *For strong internal coupling, AR shows oscillations.* ***B-F)*** *The EMT circuit shows small oscillations on top its steady states.*

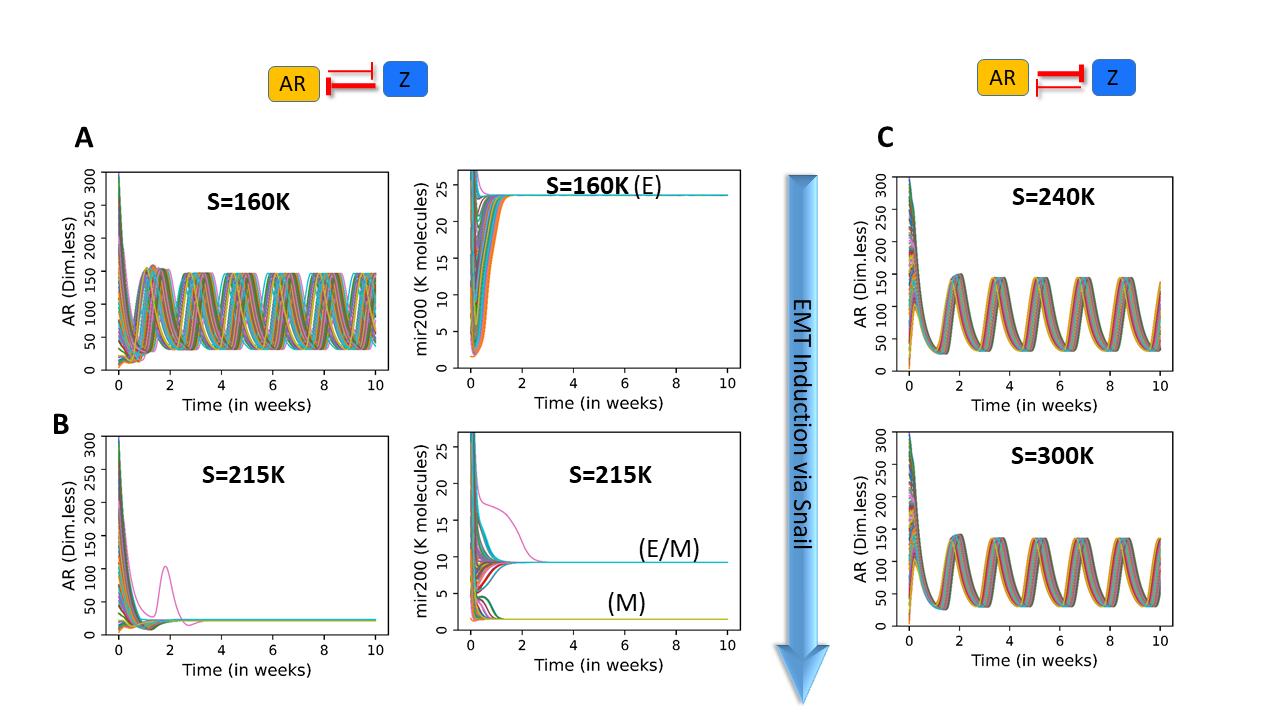

***Figure S3: Dynamics of coupled circuits. A-B)*** *Dynamics of AR and miR-200 at varied levels of SNAIL as shown in panels, and at* $\lambda_{ZtoA}$*=0.1 and* $\lambda_{AtoZ}$*=0.9:* ***C)*** *AR dynamics at* $\lambda_{ZtoA}$*=0.9 and* $\lambda_{AtoZ}$*=0.1, for varied SNAIL levels.*

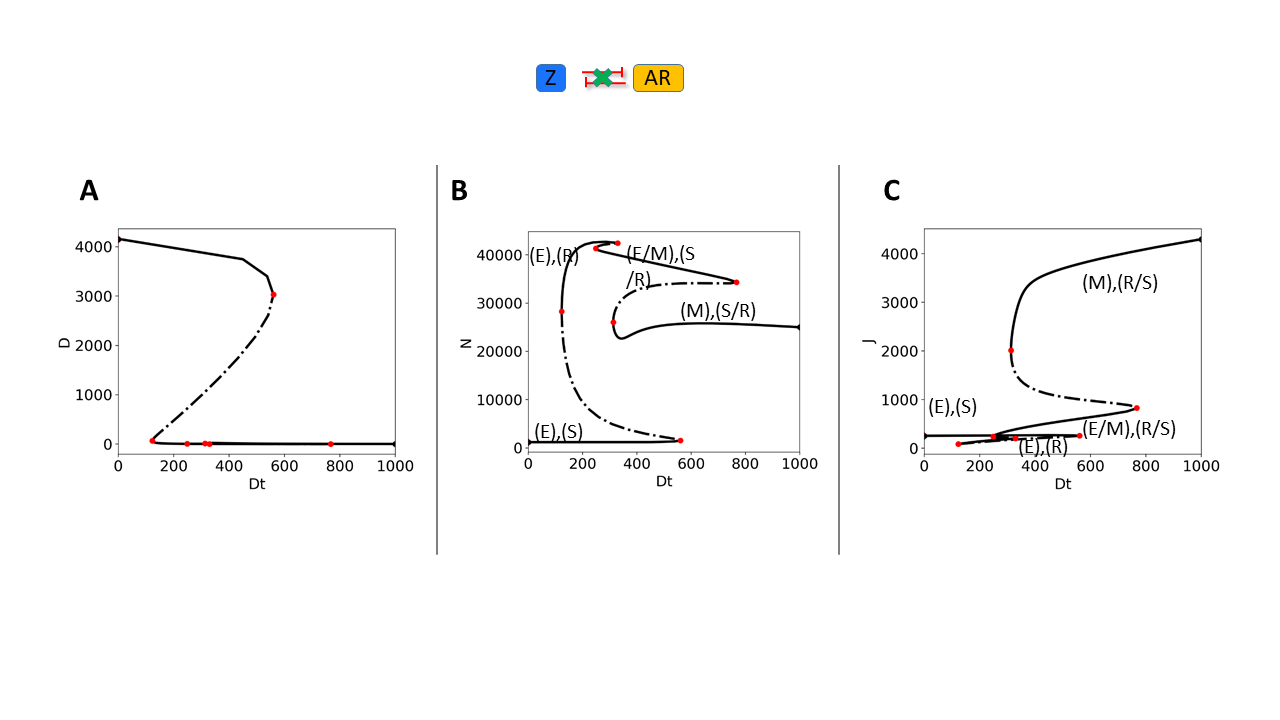
***Figure S4:*** ***Bifurcation diagram of coupled EMT-Notch signaling.*** *Bifurcation plots of Delta (D), Notch (N) and Jagged (J) with respect to External Delta (Dext, referred to as Dt in above panel) for the case when PAGE4-AR and EMT circuits are not coupled. Dt is in number of molecules.*

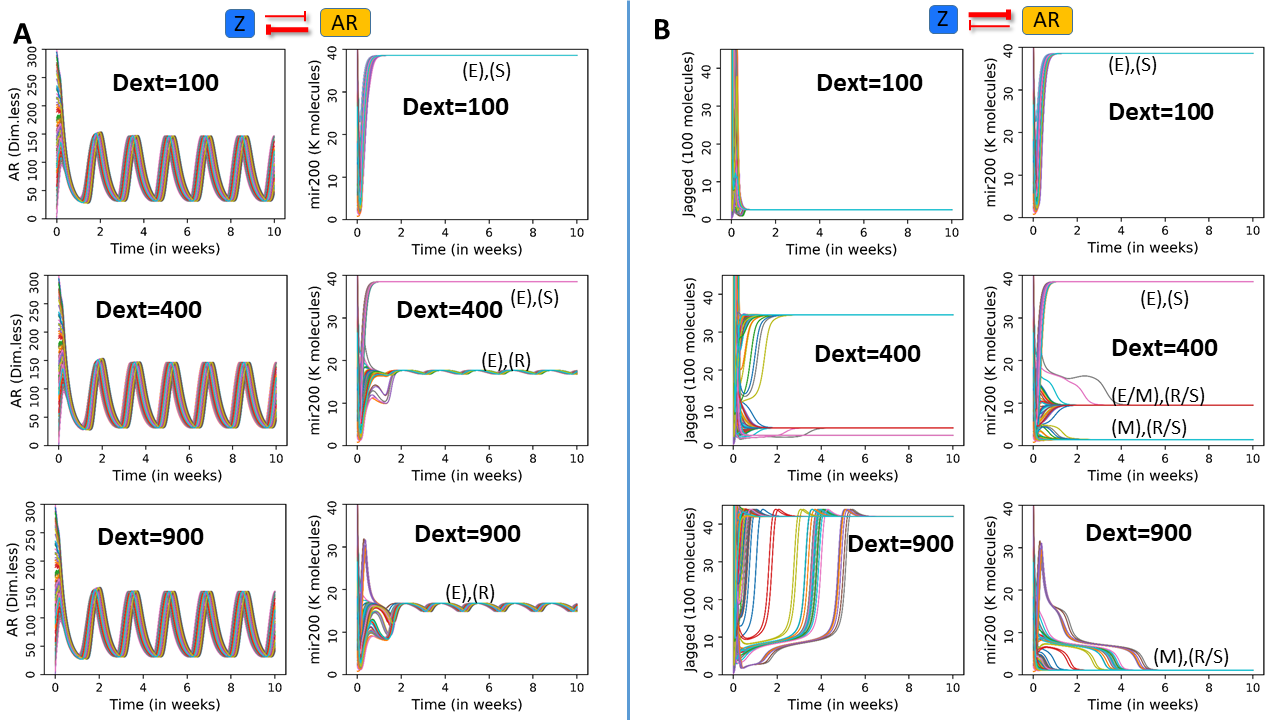

***Figure S5: Dynamics of coupled circuits at varying coupling strengths. A)*** *Trajectories of AR and miR-200 at different Snail values for the case of* $\lambda_{AtoZ}$*=0.1 and* $\lambda_{ZtoA}$*=0.9. AR continues to oscillate. For low values of External Delta (top panel), miR-200 saturates at a value in the Epithelial Sender range. At higher external Delta (middle and lower panel) the epithelial receiver state appears and shows oscillations.* ***B)*** *Trajectories of Jagged and miR-200 at different Snail values for the case of* $\lambda_{AtoZ}$*=0.9 and* $\lambda_{ZtoA}$*=0.1. Jagged and miR-200 show similar behavior as to the uncoupled circuit (Fig 5B). Dext unit is in number of molecules..*

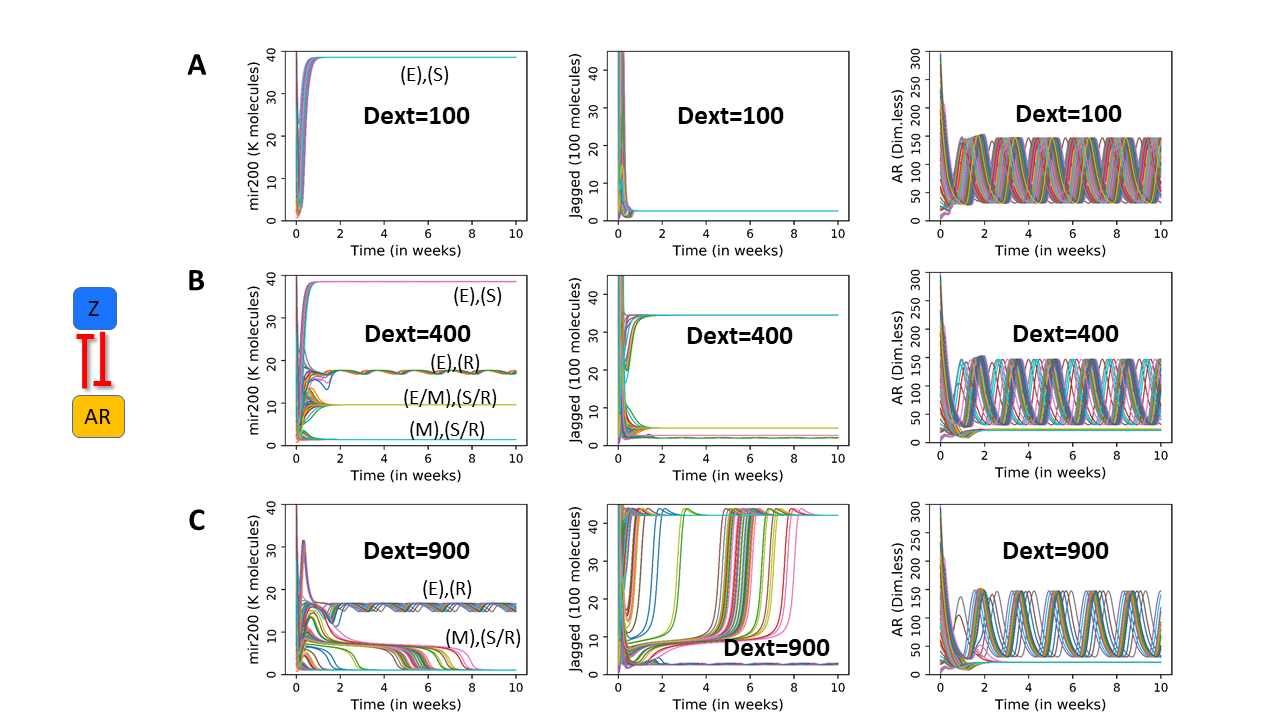
***Figure S6: PAGE4-EMT-NDJ circuit in case*** $\boldsymbol{\lambda}_{\boldsymbol{AtoZ}}$ ***and*** $\boldsymbol{\lambda}_{\boldsymbol{ZtoA}}$ ***=0.1: A)*** *On low external Delta, AR oscillates and EMT-NDJ circuit saturates at the epithelial sender state. At higher External Delta (Dext=400), cell choses either of the 4 possible states. The epithelial receiver state shows oscillations as well. At even higher External Delta (Dext=900), the hybrid and epithelial Sender state vanishes and only Epithelial Receiver and Mesenchymal (Sender/Receiver) state remain. Also, the epithelial receiver state continues to oscillates. AR either goes to an oscillatory trajectory or saturates at a low value. Dext unit is in number of molecules.*
