## Supplementary material for "Coupled feedback loops involving PAGE4, EMT and Notch signaling can give rise to non-genetic heterogeneity in prostate cancer cells": SI Materials and Methods

**Supplementary Information**

Divyoj Singh, Federico Bocci, Prakash Kulkarni, Mohit Kumar Jolly

#### 1 PAGE4 - Androgen Receptor -X Module:

Building up from our previous work in Lin et al. 2018, we first got correct dimensions of time. The absolute levels of the players are not known hence the levels are kept dimensionless. We also add another node "X" in the circuit to couple with AR.
Following are the equations for WT-PAGE4 ($P_U$), HIPK1-PAGE4 complex ($P_M$), CLK2-PAGE4 complex ($P_H$), CLK2 ($C$), Androgen Receptor ($AR$), new node "X" ($X$) :

$$\dot{P_U}(t) = g_{P_U} - H \cdot \frac{P_u(t)}{P_u(t)+a} - \gamma_{P_u} P_u(t)$$

$$\dot{P_M}(t) = H\frac{P_u(t)}{P_u(t)+a} - g_H C(t)\frac{P_M(t)}{P_M(t)+b} - \gamma_{P_M} P_M(t)$$

$$\dot{P_H}(t) = g_H C(t)\frac{P_M(t)}{P_M(t)+b} - \gamma_{P_H} P_H(t)$$

$$\dot{C}(t) = g_C H^S(A(t-\tau_C), C) - \gamma_C C(t)$$

$$\dot{A}(t) = g_A H^S(X(t), A) H^S(P_M(t-\tau_A), A) - \gamma_A A(t)$$

$$\dot{X}(t) = g_X H^S(A(t), X) H^S(X(t), X) - \gamma_X X(t)$$

The terms $H^S(X(t), A)$ and $H^S(A(t), X)$ stand for the coupling terms of AR and X. The terms $H^S(P_M(t-\tau_A), A)$ and $H^S(A(t-\tau_C), C)$ stand for the delay terms due to the intermediate terms. All parameters are taken from Lin et al. 2018 except for the parameters involving node "X" which are estimated.

Table 1: Parameters for Page4-AR circuit

| Parameter | Value | Units |
|---|---|---|
| $H$ | 2.31 | $hour^{-1}$ |
| Production rates: | | |
| $g_A$ | 4.62 | $hour^{-1}$ |
| $g_{P_U}$ | 2.31 | $hour^{-1}$ |
| $g_C$ | 2.77 | $hour^{-1}$ |
| $g_H$ | 0.04 | $hour^{-1}$ |
| $g_X$ | 2.31 | $hour^{-1}$ |
| Degradation rates: | | |
| $\gamma_C$ | 0.02 | $hour^{-1}$ |
| $\gamma_{P_M}$ | 0.004 | $hour^{-1}$ |
| $\gamma_{P_H}$ | 0.04 | $hour^{-1}$ |
| $\gamma_{P_U}$ | 0.016 | $hour^{-1}$ |
| $\gamma_A$ | 0.020 | $hour^{-1}$ |
| $\gamma_X$ | 0.04 | $hour^{-1}$ |
| Threshold constants: | | |
| $a$ | 5 | dimensionless |
| $b$ | 20 | dimensionless |
| $P0A$ | 20 | dimensionless |
| $A0$ | 65 | dimensionless |

| | | |
|---|---|---|
| $X0$ | 25 | dimensionless |
| Hill Coefficient: | | |
| $n_A$ | 4 | dimensionless |
| $n_C$ | 4 | dimensionless |
| $n_X$ | 4 | dimensionless |
| Delay constants: | | |
| $\tau_A$ | 16.23 | hour |
| $\tau_C$ | 16.23 | hour |
| Fold-change (regulation strength): | | |
| $\lambda_{AtoC}$ | 0.1 | dimensionless |
| $\lambda_{MtoA}$ | 0.1 | dimensionless |
| $\lambda_{XtoA}$ | 1 | dimensionless |
| $\lambda_{AtoX}$ | 1 | dimensionless |
| $\lambda_{XtoX}$ | 1 | dimensionless |

#### 2 PAGE4-AR-EMT Module:

To model EMT network,we first reduce the framework from Lu et al. 2013 as done in Boareto et al. 2016. Next, we replace X in the previous section with Zeb(Z). All other equations remain same. The parameters for EMT circuit are taken from Lu et al. 2013 and Boareto et al. 2016.

$$\dot{P_U}(t) = g_{P_U} - H \cdot \frac{P_u(t)}{P_u(t)+a} - \gamma_{P_u} P_u(t)$$

$$\dot{P_M}(t) = H \frac{P_u(t)}{P_u(t)+a} - g_H C(t) \frac{P_M(t)}{P_M(t)+b} - \gamma_{P_M} P_M(t)$$

$$\dot{P_H}(t) = g_H C(t) \frac{P_M(t)}{P_M(t)+b} - \gamma_{P_H} P_H(t)$$

$$\dot{A}(t) = g_A \boldsymbol{H^S(Z(t), A)} H^S(P_M(t-\tau_A), A) - \gamma_A A(t)$$

$$\dot{C}(t) = g_C H^S(A(t-\tau_C), C) - \gamma_C C(t)$$

$$\dot{Z}(t) = k_P g_Z \boldsymbol{H^S(A(t), Z)} H^S(Z(t), Z) H^S(S(t), Z) Pl(\mu_{200}(t), 6) - \gamma_Z Z(t)$$

$$\dot{\mu_{200}}(t) = g_{\mu_{200}} H^S(S(t), \mu_{200}) H^S(Z(t), \mu_{200}) - g_Z H^S(Z(t), Z) \cdot H^S(S(t), Z) Py(\mu_{200}(t), 6) - \gamma_{\mu_{200}} \mu_{200}(t)$$

Table 2: Parameters for core-EMT circuit

| **Parameter** | **Value** | **Units** |
|---|---|---|
| Translation rate: | | |
| $k_P$ | $1.0 \times 10^2$ | proteins per mRNA per hour |
| Degradation rate of mRNA: | | |
| $k_m$ | $5.0 \times 10^{-1}$ | per hour |
| Production rates: | | |
| $g_{\mu_{200}}$ | $2.1 \times 10^3$ | molecules/hour |
| $g_Z$ | 11 | molecules/hour |
| Degradation rate: | | |
| $\gamma_{\mu_{200}}$ | $5.0 \times 10^{-2}$ | $hour^{-1}$ |
| $\gamma_Z$ | $1.0 \times 10^{-1}$ | $hour^{-1}$ |
| Threshold constants: | | |
| $S0\mu_{200}$ | $1.8 \times 10^5$ | molecules |
| $S0Z$ | $1.8 \times 10^5$ | molecules |
| $Z0\mu_{200}$ | $2.2 \times 10^5$ | molecules |
| $Z0Z$ | $2.2 \times 10^5$ | molecules |
| $A0Z$ | 65 | molecules |
| $Z0A$ | $2.5 \times 10^4$ | molecules |

| Hill Coefficient: | | |
|---|---|---|
| $n_{Zto\mu_{200}}$ | 3 | dimensionless |
| $n_{Sto\mu_{200}}$ | 2 | dimensionless |
| $n_{StoZ}$ | 2 | dimensionless |
| $n_{ZtoZ}$ | 2 | dimensionless |
| $n_{ZtoA}$ | 4 | dimensionless |
| $n_{AtoZ}$ | 4 | dimensionless |
| Fold-change (regulation strength): | | |
| $\lambda_{Zto\mu_{200}}$ | 0.1 | dimensionless |
| $\lambda_{Sto\mu_{200}}$ | 0.1 | dimensionless |
| $\lambda_{StoZ}$ | 10 | dimensionless |
| $\lambda_{ZtoZ}$ | 7.5 | dimensionless |
| $\lambda_{ZtoA}$ | 0.1 | dimensionless |
| $\lambda_{AtoZ}$ | 0.1 | dimensionless |

Table 3: Parameters for translation, mRNA degradation and micro-RNA degradation upon protein-micro-RNA binding

| **Parameter group** | **Parameter** | **Value** | **Units** |
|---|---|---|---|
| Translation rate | $l_i$ | 1.0, 0.6, 0.3, 0.1, 0.05, 0.05, 0.05 | $h^{-1}$ |
| mRNA degradation rate | $\gamma_{mi}$ | 0, 0.04, 0.2, 1.0, 1.0, 1.0, 1.0 | $h^{-1}$ |
| micro-RNA degradation rate | $\gamma_{\mu i}$ | 0, 0.005, 0.05, 0.5, 0.5, 0.5, 0.5 | $h^{-1}$ |

### 3 PAGE4-EMT-NDJ:

For combining the N-D-J circuitry, we add ODE's for Snail(S), microRNA-34($\mu_{34}$), Notch (N), Delta(D), Jagged (J) and NICD(I) from Boareto et al. 2016
. All parameters are taken from Boareto et al. 2016.

$$\dot{P_U}(t) = g_{P_U} - H \cdot \frac{P_u(t)}{P_u(t)+a} - \gamma_{P_u} P_u(t)$$

$$\dot{P_M}(t) = H\frac{P_u(t)}{P_u(t)+a} - g_H C(t)\frac{P_M(t)}{P_M(t)+b} - \gamma_{P_M} P_M(t)$$

$$\dot{P_H}(t) = g_H C(t)\frac{P_M(t)}{P_M(t)+b} - \gamma_{P_H} P_H(t)$$

$$\dot{C}(t) = g_C H^S(A(t-\tau_C), C) - \gamma_C C(t)$$

$$\dot{A}(t) = g_A H^S(Z(t), A) H^S(P_M(t-\tau_A), A) - \gamma_A A(t)$$

$$\dot{Z}(t) = k_P g_Z H^S(A(t), Z) H^S(Z(t), Z) H^S(S(t), Z) Pl(\mu_{200}(t), 6) - \gamma_Z Z(t)$$

$$\dot{\mu_{200}}(t) = g_{\mu_{200}} H^S(S(t), \mu_{200}) H^S(Z(t), \mu_{200}) - g_Z H^S(Z(t), Z) \cdot H^S(S(t), Z) Py(\mu_{200}(t), 6) - g_J H^S(I(t), \mu_{200}) Py(\mu_{200}(t), 5) - \gamma_{\mu_{200}} \mu_{200}(t)$$

$$\dot{S}(t) = k_P g_S H^S(S(t), S) H^S(I(t), S) H^S(It, S) Pl(\mu_{34}(t), 2) - \gamma_S S(t)$$

$$\dot{\mu_{34}}(t) = g_{\mu_{34}} H^S(S(t), \mu_{34}) H^S(Z(t), \mu_{34}) - g_S H^S(S(t), S) H^S(I(t), S) H^S(It, S) Py(\mu_{34}(t), 2) - g_N H^S(I(t), N) Py(\mu_{34}(t), 2) - g_D H^S(I(t), D) Py(\mu_{34}(t), 3) - \gamma_{\mu_{34}} \mu_{34}(t)$$

$$\dot{N}(t) = kPg_N H^S(I(t), N) Pl(\mu_{34}(t), 2) - N(t)((kcD(t) + ktDt) H^S(I(t), D) + (kcJ(t) + ktJt) H^S(I(t), J)) - \gamma_N N(t)$$

$$\dot{D}(t) = kPg_D H^S(I(t), D) Pl(\mu_{34}(t), 3) - D(t)(kcN(t) H^S(I(t), D) + ktNt) - \gamma_D D(t)$$

$$\dot{J}(t) = kPg_J H^S(I(t), J) Pl(\mu_{200}(t), 5) - J(t)(kcN(t)H^S(I(t), J) + ktNt) - \gamma_D D(t)$$

$$\dot{I}(t) = ktN(t)(DtH^S(I(t), D) + JtH^S(I(t), J)) - \gamma_I I(t)$$

Table 4: Parameters for EMT - NDJ circuit

| **Parameter** | **Value** | **Units** |
|---|---|---|
| Production rates: | | |
| $g_{\mu_{34}}$ | $1.35 \times 10^3$ | molecules/hour |
| $g_S$ | $9 \times 10^1$ | molecules/hour |
| $g_N$ | $0.8 \times 10^1$ | molecules/hour |
| $g_D$ | $7 \times 10^1$ | molecules/hour |
| $g_J$ | $2 \times 10^1$ | molecules/hour |
| Degradation rates: | | |
| $\gamma_{\mu_{34}}$ | $5.0 \times 10^{-2}$ | $hour^{-1}$ |
| $\gamma_S$ | $1.25 \times 10^{-1}$ | $hour^{-1}$ |
| $\gamma_N$ | $1.0 \times 10^{-1}$ | $hour^{-1}$ |
| $\gamma_I$ | $5.0 \times 10^{-1}$ | $hour^{-1}$ |
| $\gamma_D$ | $1.0 \times 10^{-1}$ | $hour^{-1}$ |
| $\gamma_J$ | $1.0 \times 10^{-1}$ | $hour^{-1}$ |
| Threshold constants: | | |
| $S0\mu_{34}$ | $3 \times 10^5$ | molecules |
| $S0S$ | $2 \times 10^5$ | molecules |
| $Z0\mu_{34}$ | $2.2 \times 10^5$ | molecules |
| $I0S$ | $3 \times 10^2$ | molecules |
| $I0$ | 100 | molecules |
| Hill Coefficient: | | |
| $n_{Sto\mu_{34}}$ | 1 | dimensionless |
| $n_{StoS}$ | 1 | dimensionless |
| $n_{\mu_{34}}$ | 2 | dimensionless |
| $n_I$ | 2 | dimensionless |
| $n_F$ | 1 | dimensionless |
| $n_N, n_D, n_J$ | 2 | dimensionless |
| Fold-change (regulation strength): | | |
| $\lambda_{Sto\mu_{200}}$ | 0.1 | dimensionless |
| $\lambda_{StoS}$ | 0.1 | dimensionless |
| $\lambda_{Zto\mu_{200}}$ | 0.2 | dimensionless |
| $\lambda_{ItoS}$ | 6.5 | dimensionless |
| $\lambda_{ItoN}$ | 7.0 | dimensionless |
| $\lambda_{ItoD}$ | 0.0 | dimensionless |
| $\lambda_{ItoJ}$ | 2.0 | dimensionless |
| Cis-inhibition rate: | | |
| $kc$ | $1.0 \times 10^{-4}$ | $hour^{-1}$ |
| Trans-activation rate: | | |
| $kt$ | $1.0 \times 10^{-5}$ | $hour^{-1}$ |
| External Ligand and Notch Concentration: | | |
| $Dt$ | 0.1 | *molecules* |
| $Nt$ | 0.1 | *molecules* |
| $Jt$ | 0.1 | *molecules* |
| External Signal on Snail: | | |
| $It$ | 0.0 | *molecules* |

### 4 Functions:

In our framework, we used following functions.

$$H^S(A(t),B(t)) = H^S(A, A0B, n_{AtoB}, \lambda_{AtoB}) = \frac{1}{1+(\frac{A}{A0B})^{n_{AtoB}}} + \lambda\frac{(\frac{A}{A0B})^{n_{AtoB}}}{1+(\frac{A}{A0B})^{n_{AtoB}}}$$

We discuss Shifted Hill function in detail in Materials and Methods in the main text.

The following functions appear in the EMT circuit which models effect of micro-RNA on gene expression. The formulation is formulated in detail in Supplementary Information of Lu et al. 2013 and has been extended in Boareto et al. 2016.

$$P_l(\mu,n) = \frac{L(\mu,n)}{Y_m(\mu,n)+k_m}$$

This represents post-translational inhibition due to a micro-RNA species $\mu$ ($\mu 200 or \mu 34$ here). Here n represents the number of binding sites of micro-RNA on the promoter region of the target species. $k_m$ represents the degradation rate of mRNA.

$$P_y(\mu,n) = \frac{Y_\mu(\mu,n)}{Y_m(\mu,n)+k_m}$$

represents the decrease in the levels of microRNA due to the degradation of the microRNA/mRNA complex.
Here, $L(\mu,n) = \sum_{i=0}^{n} l_i C_i^n M_i^n(\mu)$

$Y_m(\mu,n) = \sum_{i=0}^{n} \gamma_{mi} C_i^n M_i^n(\mu)$

$Y_m(\mu,n) = \sum_{i=0}^{n} \gamma_{\mu i} C_i^n M_i^n(\mu)$

$$C_i^n = \frac{n!}{i!(n-i)!}$$

and

$$M_i^n = \frac{(\frac{\mu}{\mu 0})^i}{(1+\frac{\mu}{\mu 0})^n}$$

### 5 Phase plots:

The phase plot was divided into 50 x 50 cells. For each cell, at that parameter value, two trajectories were obtained, one starting with AR high and X low and other with AR low and X high. We used Euler-Integration with time step dt=0.001hr and for a total time of 10 weeks. After this, extrema of the second half of the trajectory were calculated, if the $\frac{difference\ of\ the\ extrema}{sum\ of\ the\ extrema} > 0.1$; then oscillatory

Otherwise, if: $\frac{differnce\ of\ steady\ state\ of\ trajectories}{sum\ of\ steady\ state\ of\ trajectories} > 0.1$; then bistable

Otherwise, it was considered monostable.
The phase plots were visualised using seaborn heatmap (Waskom and team 2020).

### 6 Dynamics Trajectories:

We first uniformly sample initial conditions from a specific range for each variable (say a random number between 0 and 300 for AR). Then we integrate the ODES's using Euler-Method with time step dt=0.001 hr and total duration of 10 weeks. The same process was repeated for 100 different trajectories with different initial conditions. The seed value was also controlled for different runs, so that results are reproducible.

### 7 Bifurcation Diagrams:

The bifurcation diagram for miR-200 in Figure 1B was made in MATLAB using the package MATCONT Dhooge, Govaerts, and Kuznetsov 2003 and was diagram is adapted from our previous study Lu et al. 2013. The bifurcation diagram for miR-200 in Figure 5B and Figure S4 were made in Python2 using PyDsTool (**ClewleyRHSherwoodWELaMarMD2007**).Figure 5B was also adapted from our previous study Boareto et al. 2016.

### 8 Codes:

These codes were written in Python3 and only basic libraries like numpy (Harris et al. 2020), scipy (Virtanen et al. 2020) and pandas (McKinney 2010) were used . The trajectories were visualised using matplotlib (Hunter 2007). All codes are available publicly on the GitHub page (https://github.com/Divyoj-Singh/PAGE4-AR-EMT-NDJ).
